# Selection for Extreme Neck Elongation in Crown Birds

**DOI:** 10.64898/2026.10.19.760280

**Authors:** Daohan Jiang, Satoshi Kobayashi, Lauren Sallan

## Abstract

Variation in segment number, length, and identity is a major axis of diversification in bilaterians, shaped by developmental novelty and constraint. The near-invariant neck vertebral (cervical) count of mammals is a classic example of such constraint, yet we find that mammals occupy an unexceptional region of amniote cervical morphospace. In contrast, crown birds evolved extraordinarily long-neck proportions through increases in both cervical number and length, a combination only approached by ecologically-distinct sauropods among other archosaurs. This condition is widespread among living birds but largely absent from flight-capable stem avialans, excluding flight as a sole driver. A model of developmental evolution of segmentation demonstrates that coordinated increases in segment number and length are a predictable response to selection for regional elongation when viable homeotic changes are accessible. Our findings show how developmental constraints shape evolutionary routes to extreme morphologies and suggest major transformations can arise in traits beyond those commonly perceived as defining a clade’s specialization.

---

Segmentation is a core driver of axial specification in the body plans of multiple bilaterian groups. Segment number can determine the absolute length of either the body or outgrowths, while differential gene expression specify the segments’ developmental identities, their resultant structures, and their positioning along the anterior-posterior axis. In vertebrates, somites become vertebrae and key musculature depending on their identities, which are determined by their placement along the axis relative to *Hox* gene expression domains [1, 2]. The evolution of segment number and identity is thus subject to both developmental and selective constraint, often displaying conservation; homeotic shifts (shifts of *Hox* gene expression domains), have also taken place and lead to evolutionary innovations in various lineages . The mammalian neck has become a classic example illustrating how developmental constraints can limit the evolution of segments; mammals almost invariably possess seven cervical vertebrae regardless of body size, proportion, or ecology, a pattern that has been attributed to strong developmental constraints associated with pleiotropy. That is, homeotic mutations that alter the number of cervical vertebrae generally have strongly deleterious pleiotropic effects and can cause severe developmental defects [3, 4]. It is in only two clades, Folivora (sloths) and Trichechidae (manatees), cervical vertebrae counts other than seven are observed, presumably as a result of homeotic shifts [5, 6].

However, very few studies have examined the inferred mammal constraint in the context of other tetrapods, limiting the explanatory power of such hypotheses. In other amniote groups, the cervical region is considerably more diverse. Sauropsids, which includes modern birds and reptiles along with all other descendants of their last common ancestor (LCA), exhibit substantial variation in cervical vertebral count as well as morphology [7]. It has been suggested that the observed diversification is likely enabled by more frequent homeotic shifts [8], indicating that the constraints operating in mammals is absent or weak. Despite this diversity, broad patterns of evolutionary divergence across amniotes yet remain incompletely characterized. Most previous studies have focused on individual clades or taxa with exceptional phenotypes (e.g., some birds [9], tanystropheids [10], sauropterygians [11]), leaving unanswered whether major amniote groups differ systematically in their modes of cervical evolution and whether these differences can be explained under a common evolutionary framework.

Gaps in comparative data for segmentation in vertebrates are accompanied by gaps in both mechanistic explanations and understanding macroevolutionary consequences. Despite extensive work in different lineages on patterns of vertebral regionalization [7, 12–16], axial elongation [17–19], and developmental processes underlying somitogenesis and regional specification [1, 8, 20, 21], it remains poorly understood how changes in segment number and length has shaped morphological diversification across vertebrate taxa. In particular, it is unclear if developmental constraints primarily limit the extent of regional elongation or determine available evolutionary outcomes under selection. Here, we address these mechanistic and empirical questions by combining a broad comparative dataset of cervical vertebral counts and relative lengths from across living and extinct amniotes with a developmental model of vertebral segmentation including somite production, *Hox*-determined regional identity, and subsequent growth. In doing so, these approaches allow us to investigate how selection and developmental constraints have interacted to shape the evolution of vertebral regions across amniotes.

To better characterize patterns and processes of cervical region evolution across major amniote groups, we conducted a broad comparative analysis of cervical region evolution across amniotes. We focused on the normalized neck length (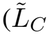, defined as neck length divided by the total length of thoracic and sacral regions of the vertebral column), as well as the cervical vertebrae count (*n*_*C*_) (Fig. 1A). The first comparison recovered a rough morphospace based on these two axes of variation (Fig. 1B). Neck lengths of mammals fall roughly in line with expectations from amniotes with equivalent numbers of cervical vertebrae, and 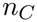 is generally correlated with *n*_*C*_ among amniotes. However, two groups of dinosaurs do show dramatic shifts in terms of linked vertebral count and overall neck lengths: the sauropodomorphs and crown birds (LCA of modern birds and all its descendants).

**Figure 1:**
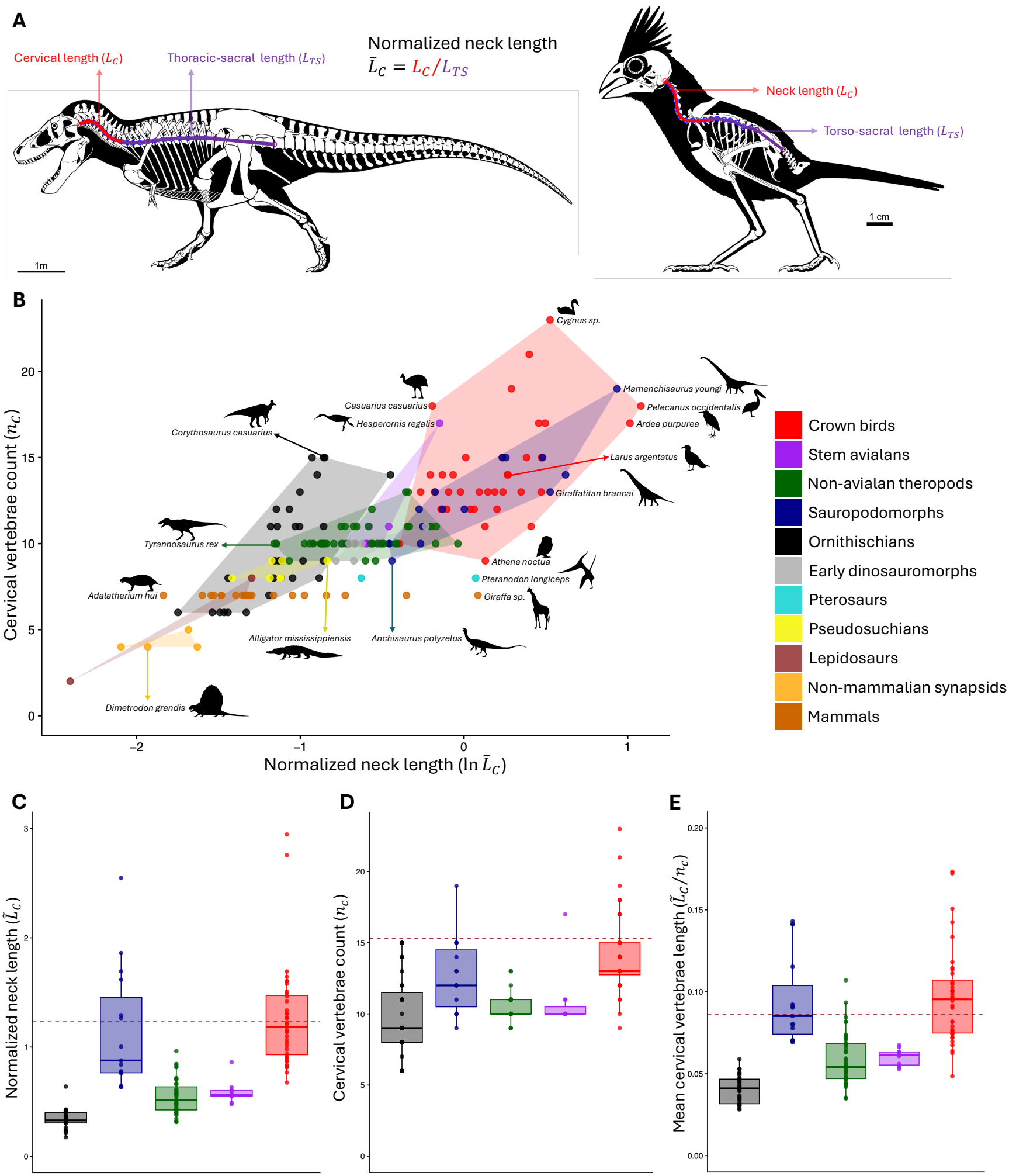
Crown birds show exceptional neck elongation relative to most other amniotes, including other archosaurs. (A) Definition of the primary phenotype of interest, with skeletal drawings of a *Tyrannosaurus rex* and a northern cardinal (*Cardinalis cardinalis*, wing removed for visual clarity) used as examples. We measured the length of the cervical region (*L*_*C*_, length of the red curve) and the total length of thoracic and sacral regions (*L*_*T S*_, length of the purple curve). The normalized neck length is calculated as 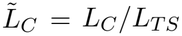. Modified from Scott Hartman. (B) Log normalized neck lengths 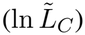 and cervical vertebrae counts (*n*_*C*_) across groups. Phenotypic subspaces occupied by several groups are visualized as polygons of matching colors. Some representative species are highlighted. Silhouettes were downloaded from PhyloPic. (C-E) Distribution of cervical phenotypes among dinosaur groups. (C) Normalized neck length 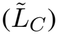. (D) Cervical vertebral count (*n*_*C*_). (E) Mean cervical vertebrae length, calculated as 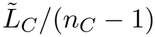 (the atlas is excluded). Dotted lines in (C-E) represent estimated ancestral states of crown birds. “Crown birds” refers to descendants of LCA of all modern birds. “Avialans” refers to members of clade Avialae, which includes descendants of LCA of crown birds, *Archaeopteryx*, and *Anchiornis*; “stem avialans” refers to avialans that are not crown birds. “Early dinosauromorphs” refers to taxa that are closer to theropods, sauropodomorphs, and ornithischians than to pterosaurs but do not belong to or cannot be confidently assigned to any one of the three groups. “Pseudosuchians” refers to archosaurs that are more closely related to crocodilians than to birds. Definitions for the clades referred to in (B-E) are listed in Table S1.

Indeed, all crown birds occupy an area of morphospace that is markedly different from all other archosaurs except sauropods; closer relatives of crown birds, such as dromaeosaurids, troodontids, and flightcapable stem avalians like Enantornithines, do not generally possess exceedingly long necks (Fig. 1B). Differences between dinosaur groups in neck length, cervical vertebrae count, and cervical vertebrae length show concordance: birds generally have proportionally longer necks (Fig. 1C), more cervical vertebrae (Fig. 1D), and longer cervical vertebrae (Fig. 1E). The inferred values of these traits in the common ancestor of crown birds exceed most other archosaurs apart from a few outliers among sauropods, or crown-ward birds (Fig. 1C-E), indicating the differences across groups cannot be explained by an over-representation of bird species from specific clades.

To investigate the timing of the evolutionary shift leading to the distinct distribution observed in crown birds, we reconstructed ancestral states for 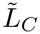. We considered three alternative topologies for Paraves phylogeny (Fig. S1) and generated 100 randomly node-dated trees for each topology, yielding 300 trees that are consistent with known tip ages. We repeated the analysis across all 300 trees to account for uncertainty in both phylogenetic relationships and divergence times. We found shift toward higher 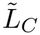 near the origin of the crown group, specifically in the stem branch leading to Euornithes (including crown birds and *Hesper-ornis*) and that leading to crown birds (Fig. 2A). The inferred ancestral value 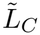 was high at the ancestral node of crown birds and moderately high at the ancestral node of Euornithes (Fig. 2B). In contrast, trait values reconstructed at older ancestral nodes, including those of Ornithothoraces, Pygostylia, Avialae, Paraves, Maniraptora, Coelurosauria, and Tetanurae, were not particularly high (Fig. 2B), though the similarity between ancestral nodes of Avialae, Paraves, and Maniraptora may in part be attributable to their close ages. Analyses using different Paraves tree topologies and different methods of ancestral state reconstruction recovered similar trends (Fig. S2). The average length of cervical vertebrae, quantified as 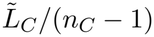, showed a similar increase after ancestors of Euornithes and Enatiornithes split (Fig. S3). These results suggest that the evolutionary shift toward the distinct distribution of 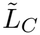 characteristic of crown birds occurred relatively late in avian evolution, after the Euornithes-Enatiornithes split.

**Figure 2:**
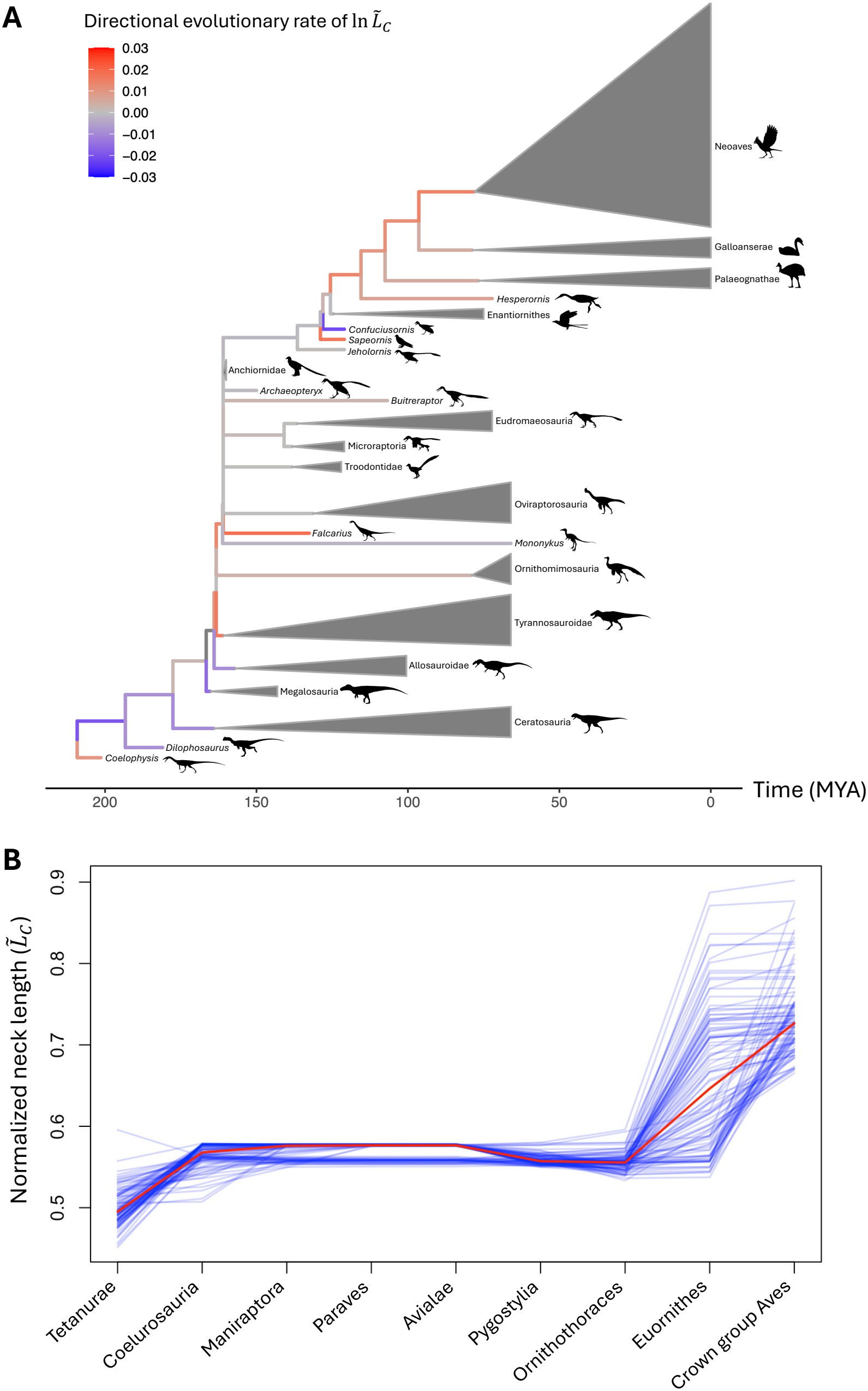
Evolution of neck length in theropods. (A) Backbone phylogeny of theropods with branches colored according to the inferred directional evolutionary rate (net evolutionary change divided by branch length) of log normalized neck length 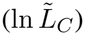. The analysis was repeated across 100 randomly generated node-dated trees sharing the same topology and tip-dating; the tree shown is an example presented for visualization. (B) Inferred ancestral states of 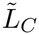 of clades that include crown birds. From left to right are progressively younger clades. Each blue curve corresponds to a random node-dated tree, while the red curves shows the median. Definitions of the clades are listed in Table S1. Results shown in this figure are based on the extended implied weighting tree reported by Motta et al. (2025) [49].

To explain the observed patterns of evolutionary divergence and explore the selective and developmental mechanisms underlying the unique patterns seen in birds, we formulated a model for the evolution of segmentation that can be applied to vertebrae as well as other segment-like structures. Here, the number of segments in each body region is determined by the expression domain of the corresponding homeotic gene (Fig. 3A), and the length of a given region can be altered by two types of mutations, which affect stages of patterning and later growth processes, respectively [22]. Homeotic mutations that cause the expression domain of the a homeotic gene to expand or shrink can alter the number of segments in the corresponding region (Fig. 3B), whereas mutations that affects growth processes regulated by the homeotic genes modify lengths of individual segments (Fig. 3C). While both types of mutation can potentially contribute to response to selection on the total length, the course of evolution will be contingent on the availability of these two types of mutations, as homeotic mutations are usually pleiotropic and strongly deleterious such that the effective rate of non-lethal homeotic mutations very low [23].

**Figure 3:**
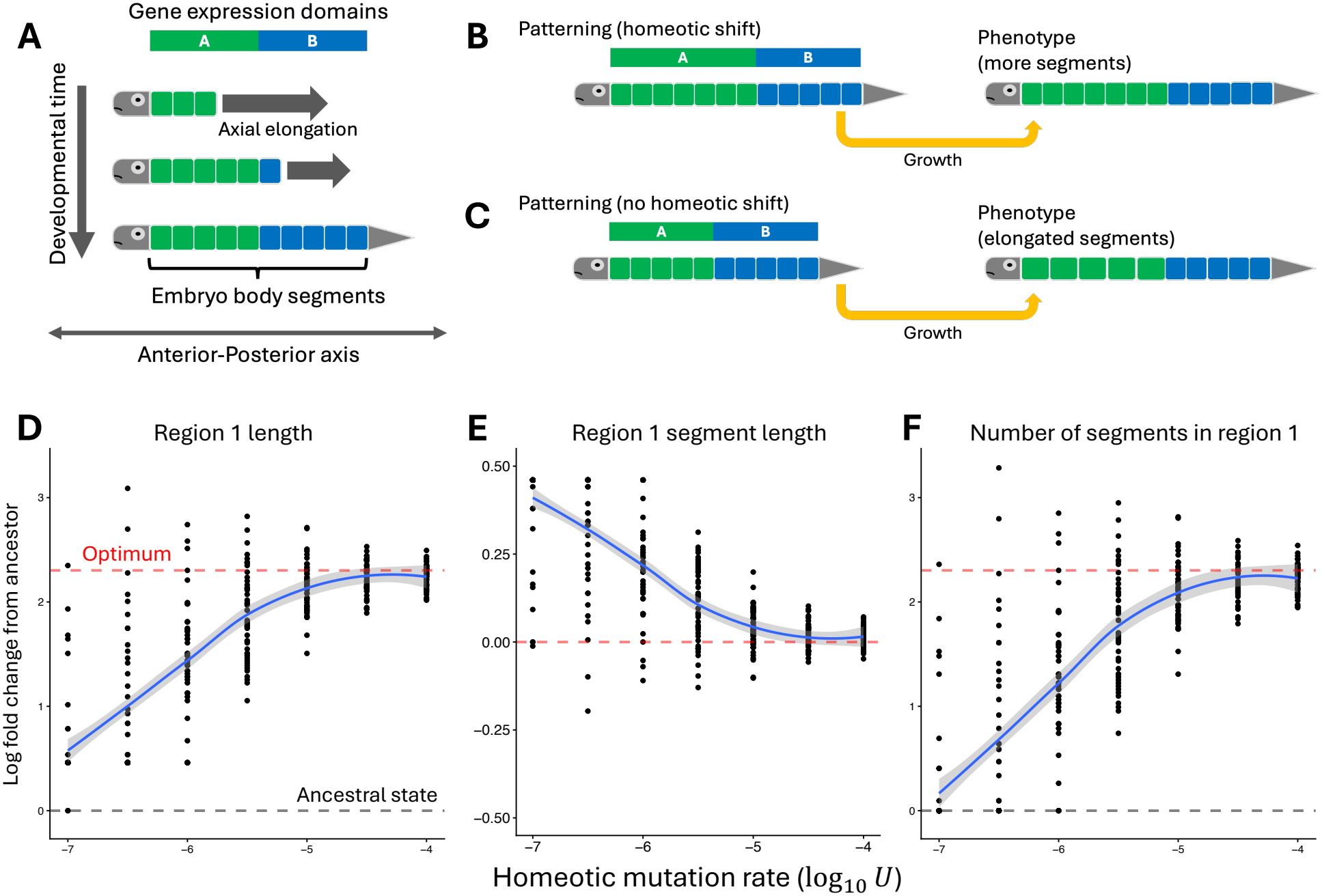
Simulations of segmentation evolution under selection for neck elongation. (A) Schematic illustration for the developmental model of segmentation, where embryonic segments are added sequentially into expression domains of the homeotic genes during development. Embryonic segments in expression domains of different homeotic genes will grow into segments comprising the respective body regions. (B-C) Schematic illustrations of genetic changes that result in the elongation of the anterior-most region (region 1). (B) Expression domain of the homeotic gene that specifies region 1 identity is expanded (a homeotic shift) such that region 1 has more segments. (C) Individual segments in region 1 are elongated due to changes in post-patterning growth processes. (D-F) Response to selection in the evolutionary simulations. The total length of region 1 was under directional selection while individual segment lengths are under stabilizing selection. Each data point is the phenotypic state of a replicate lineage at the end of the simulation. All traits were normalized and represented as log fold change from the ancestor. (D) The total length of region 1. The gray dashed line is the ancestral length and the red dashed line is the optimal length. (E) Individual segment length of region 1. The red dashed line is the optimum if the total length of the region is not under selection. (F) The number of segments in region 1. The gray dashed lines represent ancestral states and the red dashed line is the optimal number if individual segment length cannot change. Blue curves in (D-F) are the locally estimated scatterplot smoothing (LOESS) curves and the gray shaded areas are 95% CIs.

Using evolutionary simulations under the model, we explored the dynamics of evolution under a range of parameter combinations. We find the general patterns seen in the presence of directional selection on the total length of an anterior region and stabilizing selection on individual segment lengths is consistent with the observations in crown birds. Response to selection for regional elongation usually leads to concordant, complementary evolutionary changes in the number of segments in the selected region and the length of individual segments (Fig. 3D-E, Fig. S4-S6). When the rate of non-lethal homeotic mutations is low, evolution often ended up in sub-optimal states where individual segment lengths are extreme (Fig. 3D-E, Fig. S4-S6). Such scenarios are reminiscent of the cervical region of mammals, where the number of vertebrae is strongly constrained due to pleiotropy [5], and so extremely long necks are produced via elongation rather than addition of neck vertebrae rather (e.g., the giraffe). As the rate of such mutations increases, response to directional selection on region length becomes increasingly mediated by evolutionary changes in the number of segments and less by changes in segment lengths, especially when stabilizing selection on individual segment length is strong (Fig. 3D-E, Fig. Fig. S4-S6). Taken together, our model show the neck elongation phylogenetically and temporarily coincident with the appearance of the origin of crown birds can be explained by selection on the total length and moderate availability of homeotic mutations.

Together, our results suggest crown birds occupy a novel adaptive zone (a phenotypic subspace conferring high fitness) or macroevolutionary landscape (i.e., a phenotypic subspace where the adaptive peak can move) (Fig. 4A). The extreme neck elongation involved increases in both cervical vertebrae number and length, consistent with concordant homeotic and non-homeotic changes predicted by our model. Sauropods, which evolved long necks in association with large-bodied herbivory, underwent a similar, convergent shift, though their vertebral counts did not reach the highest values observed in some living birds. Mammals, in contrast, are generally confined in a lower-dimensional subspace characterized by a fixed number of cervical vertebrae and neck lengths that are unexceptional for amniotes, with the extreme neck elongation in giraffes achieved via non-homeotic changes for reasons similar to those of sauropods.

**Figure 4:**
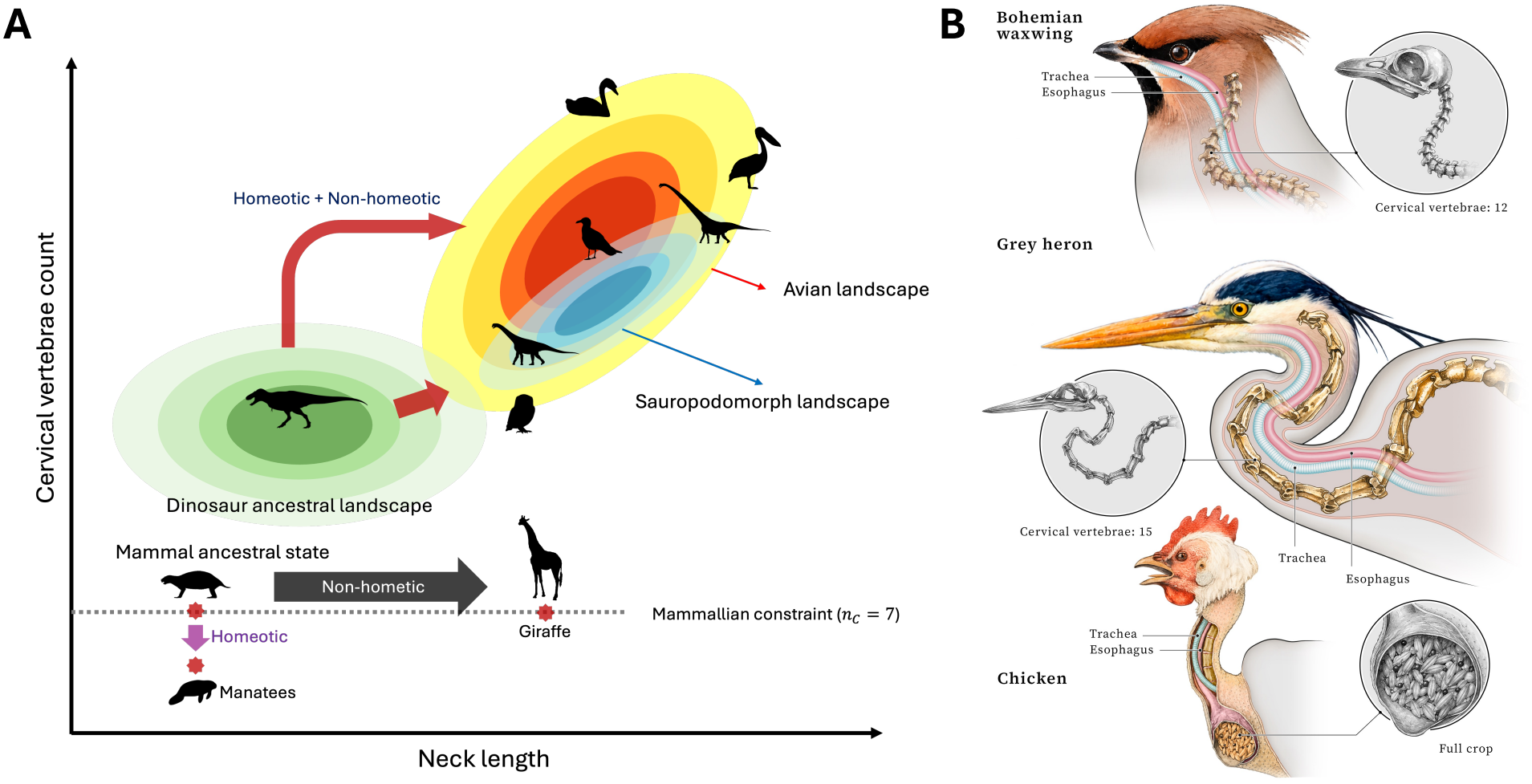
Macroevolutionary landscape for neck length and vertebral counts and examples of elongated neck in crown birds. (A) Schematic illustration of an macroevolutionary landscape for neck length and cervical vertebrae count in dinosaurs and mammals. Mammals mainly occupy a one-dimensional phenotypic subspace of *n*_*C*_ = 7, though some groups (sloths and manatees) deviated from this line via homeotic shifts. Giraffes evolved long neck via the elongation of cervical vertebrae. In contrast, birds and sauropods evolved long necks via a combination of homeotic and non-homeotic shifts, with birds more extreme values of *n*_*C*_ reached in crown birds. (B) Neck anatomy of three representative bird species, showing the cervical spine, the esophagus, and the trachea. A Bohemian waxwing (*Bombycilla garrulus*), a grey heron (*Ardea cinerea*), and a chicken (*Gallus gallus*) are used as examples. Illustrations of different species are shown at different scales. Illustration by Karin Spijker.

What selective pressures could have produced the exceptional neck elongation of birds? Flight is usually the default assumption for modifications in this clade, as a divergent lifestyle would impose novel functional demands and constraints. However, this is not a sufficient explanation; flight-capable stem avialans like the enantiornithines did not generally possess exceptionally long necks, suggesting that flight alone does not impose strong selection for neck elongation. Conversely, the stem avialan in our dataset with the highest cervical vertebrae count, the Hesperornis, was closely related to crown birds yet flightless [24]. Likewise, reduction of the forelimbs has been proposed as a driver of increased neck flexibility, which is in turn enabled by increased length [25]. However, in addition to the flighted examples noted above, several non-avian theropod lineages underwent convergent reduction of the forelimbs [26] without evolving unusually long necks, and even in some cases falling at the lower end of observed lengths (tyrannosaurids and carcharodontosaurids). Thus, neither flight nor forelimb reduction requires neck elongation as a solution.

Nevertheless, neck elongation may have been enabled or required by flight-related or other earlier modifications to the evolving avian body plan. Birds show reduction and reorganization in muscular connections between the neck, pectoral girdle and skull, with some requisite changes occurring within earlier theropods [27, 28]. These changes may have relaxed functional constraints on the neck, allowing it to evolve as a semi-independent module. Birds also possess heavily modified neck musculature relative to other living sauropsids, which allows the esophagus (along with the trachea) to be asymmetrically situated along the spinal column, creating a compartment which enables easier extension and retraction without disrupting feeding or breathing [27] (Fig 4B). The neck itself is more flexible than in other amniotes, with three distinct regions of motion approximating the S-curve [29]. Finally, an extended anterior compartment may have compensated for space limitations imposed on the chest by enlarged flight musculature and pectoral elements and an extensive air sac system. Indeed, many digestive and food storage functions have been moved to the elongate esophagus enabled by neck extension in modern birds [30–32]. This may contribute to the maintenance of relative long neck length even in birds without apparent mobility needs (Fig. 4B). Regardless of the initial drivers, the long neck was co-opted and further modified in adaptations to different ecological niches and feeding strategies as birds diversified [33, 33, 34], and so may have provided a new axis of divergence.

While the origin of birds entailed a variety of evolutionary innovations, many of the novelties, such as feathers [35–37], pygostyle [38–40], and powered flight [41, 42], are not unique to crown birds but originated well before they split from their relatives. The extreme neck elongation, in contrast, distinguishes living birds from their close relatives like Enantornithines. Such an innovation could have altered competitive relationships within an otherwise saturated ecological landscape, potentially contributing to turnover [43]. Alternatively, advantages in feeding efficiency, resource storage, or functional versatility may have influenced extinction selectivity, including the differential survival of lineages across the K-Pg boundary and afterwards [44, 45].

Our findings reveal the avian neck as an evolutionary innovation, achieved via by the coordinated evolution of cervical vertebrae count and morphology. This shows that living clades can accumulate distinct innovations at different points in their evolutionary history, either enabled by their initial trait syndrome (flight in this case) or initially incidental to it but enabling refinements in life mode or new axes and avenues of variation in saturated spaces. Finally, The contrast between birds, non-avian dinosaurs, and mammals illustrates how the interplay of developmental constraint and selection can shape alternative evolutionary responses to selection on a body region.

## Materials and Methods

### Morphometric data

We measured cervical, thoracic and sacral region lengths using skeletal images of 177 extant and extinct amniote species where complete and verified vertebral count information was available. Fossil taxa that are closely related to extant archosaurs and mammals are included in our dataset. Specifically, fossil taxa in our dataset include dinosaurs, pterosaurs, pseudosuchians (crocodilians and archosaurs that are more closely related to crocodilians than to birds), and synapsids (mammals and amniotes that are more closely related to mammals than to birds) are included in our dataset. Obligatory aquatic species (e.g., cetaceans) were not included for being too ecologically distinct. Cingulatas, ankylosaurids, and some ceratopsids were excluded as in most image data their cervical vertebrae are too severely obscured by other bony elements. These data were not widely available in existing bird compendia, perhaps because most museum specimens consist of skins without internal skeletons, and because of the fragility of the bones. Taxa included in our analysis and the data source for each are listed in Data S1.

For each taxon, the first thoracic vertebra was identified as a vertebra near the shoulder girdle that is morphologically distinct from all of the more anterior vertebrae. Vertebrae anterior to the first thoracic vertebrae are defined as cervical. We also referred to counts reported by Böhmer et al. (2019) to crossverify our classification [9]. The neck length (*L*_*C*_) is defined as the length along centra from the first cervical vertebra that is visible in the lateral view to the last cervical vertebrae. For sauropsids, the atlas is generally invisible in the lateral view and included in neck length. The torso-sacral length is defined as the length along centra from the first thoracic vertebrae to the last sacral vertebrae. The normalized neck length 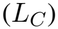 is defined as the neck length divided by the torso-sacral length. The mean cervical vertebrae length was calculated by dividing the normalized by the number of visible cervical vertebrae, which would be 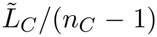 in sauropsids and 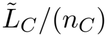 for synapsids.

Image digitization was done with R package geomorph [46]. A landmark was placed at the anterior end of the centrum of each cervical and thoracic vertebra. For the sacral region (including vertebrae fused into the synsacrum in birds), one landmark is placed at the anterior end and another is place at its posterior end. Landmark coordinates were extracted from the resulting shape files, and the lengths of cervical and torso-sacral curves were calculated as the sum of Euclidean distances between consecutive landmarks along each curve (Fig. 1A).

### Phylogenetic analysis of modern birds

A phylogenetic tree of 29 crown bird species were obtained from TimeTree5 [47]. The log normalized neck length 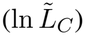 and the log mean length of visible cervical vertebrae 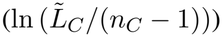 were fitted to models of continuous trait evolution using the fitContinuous function of R package geiger [48]. The estimate of trait values at the root under the best-fit model was used to represent the ancestral states of crown birds. An ancestral state estimate for *n*_*C*_ *−* 1 was calculated by dividing the ancestral state estimate of 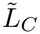 by the ancestral state estimate of 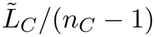. To verify the result of this analysis as well as phylogenetic representativeness of our dataset, we analyzed a dataset of bird cervical vertebrae counts that includes more species [9]. We analyzed ln *n*_*C*_ *−* 1 of 91 extant bird species, using the same method, estimated the ancestral value, and compared the corresponding ancestral *n*_*C*_ to that obtained from two datasets.

### Phylogenetic analysis with fossil taxa

For the phylogeny of Paraves (descendants of LCA of modern birds, *Archaeopteryx*, and *Anchiornis*), we considered two alternative topologies presented in Motta et al. (2025) [49] and a topology from Pei et al. (2020) [41]. In the topologies presented by Motta et al., Unenlagiidae (represented by one species, *Buitreraptor gonzalezorum*, in this study) is outside Dromaeosauridae and is closer to Paraves. In one of the topologies, *Archaeopteryx* and Anchiornithidae (represented by three species in this study: *Archiornis huxleyi, Aurornis xui*, and *Xiaotingia zhengi*) are closer to each other than to other members of Avialae; in the other tolopogy, Anchiornithidae is the earliest-diverging Avialae lineage. In Pei et al.’s topology, Unenlagiidae is within Dromaeosauridae and Anchiornithidae is the earliest-diverging Avialae lineage. The Tyrannosauroidae tree topology follows [50]. Topology of the rest of the tree follows Hendrickx et al. (2015) [51]. Non-Paravian theropods whose phylogenetic position cannot be confidently settled within this framework were excluded from the phylogenetic analysis; *Tanycolagreus topwilsoni, Monolophosaurus jiangi, Neovenator salerii, Suchomimus tenerensis*, and members of Compsognathidae were thus excluded.

Dated trees were obtained using the `cal3TimePaleoPhy` function of R package `paleotree`, which generates random dated trees given the tree topology with node ages constrained by known tip ages [52]. Tip ages of fossil taxa used for this study are listed in Data S2. Following Benson et al. (2018), speciation and extinction rates were set to be equal and the sampling rate were set lower than speciation and extinction rates [53]. For each of the three topologies, we obtained 100 random dated trees. We compared results of analysis using these different trees to see if the conclusions are robust. Ancestral reconstruction were performed for ln 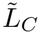 and ln 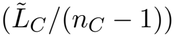 using the `RRphylo` function of the `RRphylo` package [54] and the fastAnc function of phytools [55]. We examined the ancestral states of multiple clades that contain crown birds, including Euornithes, Ornithothoraces, Pygostylia, Avialae, Paraves, Maniraptora, Coelurosauria, and Tetanurae. Definitions of these clades follow established phylogenetic nomenclature, but their operational definitions are specified (listed in Table S1) to ensure consistent taxonomic assignment across the phylogenetic tree used in this study.

### Model of developmental evolution

We considered a set of embryonic segments, which will eventually grow into a set of adult segments (e.g., vertebrae), laid along an anterior-posterior axis. Identities (developmental fates) of the segments are specified by the homeotic genes, which encode *trans*-regulatory factors. Each homeotic gene has an expression domain along the embryonic anterior-posterior axis. Spatial ranges of the embryonic segments and expression domains of the homeotic genes can be described by a coordinate system where the anterior end of the anterior-most segment is 0.

In this study, we focused on a model of vertebrate segmentation that is based on segmentation processes regulated via the *Hox* timer and the somitic clock [56]. Under this model, embryonic segments (somites) are sequentially added to the posterior end during the segmentation proceeds, until the there is no homeotic gene expressed in the corresponding location. The spatial range of the *i*-th embryonic segment (counting from the anterior end) is designated as [*i −* 1, *i*). For simplicity, we assumed that the homeotic genes’ expression domains are non-overlapping, with size of the *i*-th expression domain denoted as *D*_*i*_ and the span of the domain equal to 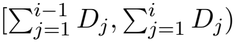. The number of embryonic segments in the *i*-th expression domain is a ceiling function of *D*_*i*_ *n*_*i*_ = ⌈*D*_*i*_⌉ . Adult segments derived from embryonic segments in the *i*-th homeotic gene’s expression domain together form the *i*-th body region (e.g., a region of the vertebral column). When *D*_*i*_ is altered due to a mutation and becomes *D*_*i*_ + *δD* (*δD > − D*_*i*_), expression domains that are posterior to the *i*-th expression domain will move accordingly, with both ends of each domain moving posterior-ward by |*δD*| when *δD >* 0 and move anterior-ward by |*δD*| when *δD <* 0.

After segmentation is completed and identities of the segments are settled, growth takes place and transform the embryonic segments into adult segments. The phenotypic state of each adult segment is determined by the expression of the effector genes during growth. We assumed that the expression of the effector genes in a growing segment is regulated by the homeotic gene expressed at the anterior end of the corresponding embryonic segment. When there are *p* homeotic genes and *q* effector genes, the phenotypic state of a given adult segment, represented by a vector 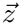, is given by

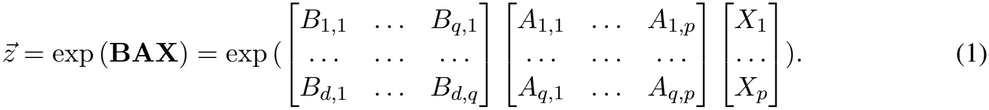

Here, **X** is a column vector containing local expression states of the homeotic gene. In this study, it is assumed that the expression domains of different homeotic genes do not overlap, so there will be one element of **X** that is equal to one and the rest will be zeros. The matrix **A** captures regulatory effects of the homeotic genes on the effector genes, with element *A*_*i,j*_ representing the *j*-th homeotic gene’s per-unit-expression effect on the *i*-th effector gene. The matrix **B** characterizes each effector gene’s effects on the phenotypic state, with element *B*_*i,j*_ representing the *j*-th effector gene’s per-unit-expression effect on the *i*-th phenotypic dimension (trait). The exponential conversion makes elements of 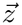 non-negative and appropriate for representing organismal traits like length, volume, or mass of a body part.

For simplicity, in this study, we focused on a simple case where the phenotypic state of interest is one-dimensional and represented by a scalar *z*. Together, for the *i*-th body region, Eqn. (1) can be written as

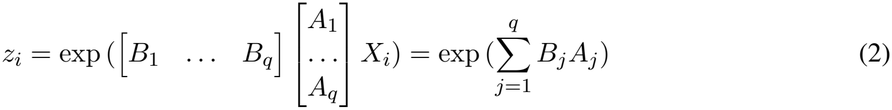

In this study, we interpret *z*_*i*_ as the length of each adult segment in the *i*-th region. The total length of the *i*-th body region is then *n*_*i*_*z*_*i*_, where *n*_*i*_ is the number of segment-derived parts in this region.

It can be seen from Eqn. (2) that the log-transformation of *z*_*i*_ is a sum of monomials that reflect the effector genes’ phenotypic effects and their response to the *i*-th homeotic gene. Assuming that mutations’ effects on these terms are generally small and additive, the evolution ln *z*_*i*_ can be modeled as a quantitative trait with an infinitesimal genetic architecture and readily approximated by BM or OU processes.

Phenotypic traits under selection include the length of each adult segment in each body region (summarized by a vector **z** = [*z*_1_ … *z*_*i*_] ^*⊤*^ if there are *i* regions) and the total length of each region (summarized by a vector **Z** = **n** ◦ **z** =[*n*_1_*z*_1_ … *n*_*i*_*z*_*i*_] ^*⊤*^ if there are *i* regions). Fitness with respect to **Z** is a multivariate Gaussian function of ln **Z**:

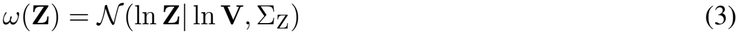

where vector **V** is the optimum of **Z** and Σ_*Z*_ captures the shape of the fitness function.

Fitness with respect to **z** is a product of two components. The first fitness component is independent of **Z** and is a multivariate Gaussian function of ln **z**:

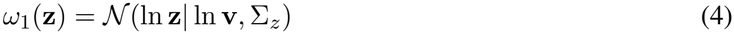

where **v** is the optimum of **z** when **Z** is not under selection and Σ_*z*_ characterizes the shape of the fitness function. In this study, we considered scenarios where Σ_*z*_ is diagonal and all diagonal elements of Σ_*z*_ are equal. The second component, which is mediated by selection on **Z**, is given by

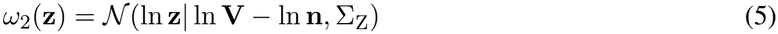

Together, the effective fitness function for ln **z** is

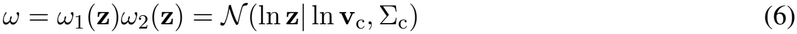

where

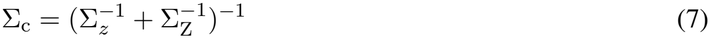

and

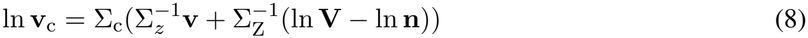

Given **n**, the evolution of ln **z** can be modeled as an OU process where the instantaneous rate of evolution at time *t* is:

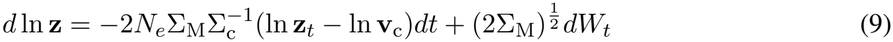

where where *N*_*e*_ is the effective population size and Σ_M_ is the mutational variance-covariance matrix for segment lengths (per haploid genome), with each row (column) corresponding to a region. Neutral evolutionary change in ln **n** over time *T* thus follows a multivariate normal distribution *N* (**0**, 2Σ_M_*T* ). Expected evolutionary change due to selection given the ancestral state **z**_0_ is equal to 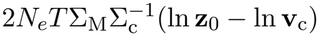.

### Evolutionary simulations

It is assumed that mutations that affect expression domains of the homeotic genes are usually highly pleiotropic and thus strongly deleterious. After excluding mutations that are make reproductive fitness zero, the effective rate of mutations that affect sizes of homeotic gene expression domains (homeotic mutations for short) is low enough to be compatible with a sequential fixation model [57]. To simulate evolution over time *T* in a lineage, we first sampled the number of homeotic mutations that would occur during the simulation from a Poisson distribution with mean equal to 2*N*_*e*_*UpT, U* is the effective homeotic mutation rate per homeotic gene per haploid genome, and *p* is the number of homeotic genes. We then determined time at which each mutation occur by sampling from a uniform distribution U(0, *T* ).

Evolutionary change of **z** between homeotic mutations was modeled by Eqn. (9). If the time interval between between a given homeotic mutation (e.g., the *t*-th homeotic mutation) and the previous homeotic mutation (e.g., the *t −* 1-th homeotic mutation) is Δ*t*, the total amount of neutral evolutionary change Δ ln **z**_neutral_ would first be sampled from a multivariate normal distribution *N*(**0**, 2Σ_*M*_ *T* ). We then computed the evolutionary change of ln **z** due to selection as 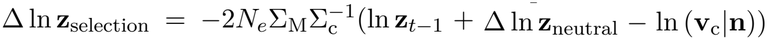, where **z**_*t* 1_ is the ancestral state at the beginning of the time interval after the *t −* 1-th homeotic mutation was either fixed or purged by selection, and ln (**v**_c_ **n**) is the effective optimum of **z** given **n**. Together, before the *t*-th homeotic mutation, there is ln **z**_*t*_ = ln **z**_*t−*1_ +Δ ln **z**_neutral_ +Δ ln **z**_*selection*_. After obtaining ln **z**_*t*_, we calculated fitness effect of the focal homeotic mutation. To determine its effect, we first randomly chose a homeotic gene whose expression domain would be affected by the focal mutation. If *D*_*i*_ is affected, the mutant value will be *D*_*i*_ exp (*δ*), where *δ* is sampled from a normal distribution *N*(0, 1). Mutant fitness with respect to **Z** (and *x*, if applicable) were then calculated based on expression domains in the mutant. In simulations where *x* is under selection, the ancestral value of *D*_1_ was set as the optimum of *x*. We considered two homeotic genes whose ancestral expression domains are [0, 10) and [10, 20), respectively, which means there is **n** = [10 10] ^*⊤*^ ancestrally. The ancestral state and optimum of **z** was **z** = [1 1]^*⊤*^. The ancestral value of **Z** is thus **Z** = [10 10]^*⊤*^.

We considered scenarios where *Z*_1_ is under directional selection for a greater value while *Z*_2_ is under stabilizing selection. For our simulations, we considered optima of *Z*_1_ that are 2, 4, 6, 8, and 10 times of the ancestral values. In our simulations, diagonal elements of Σ_*z*_ were all equal while off-diagonal elements are zeros. Values considered for diagonal elements of Σ_*z*_ include 1.25, 2, and 5, which we referred to as representing “strong”, “moderate”, and “weak” stabilizing selection on segment lengths. Parameter combinations that were used in the simulations and the corresponding outcomes are listed in Data S3.

All analyses were conducted in an R environment [58], unless specified otherwise. Licensing information for the animal silhouettes used in the figures are summarized in Data S4.

## Supplementary text

### Limitation of sampling

Our morphometric dataset is subject to several limitations in terms of species sampling. The first major limitation comes from the fact that museum specimens of birds are most commonly preserved as study skins, during the preparation of which the much of the vertebral column is lost [59–61]. Skeletal collections, in comparison, is much less substantial, and suffers from taxonomic bias [62]. It is not clear how this bias propagate into the subset of specimens that are digitized and available to the present study. Yet, some highly diverse groups are not represented by a proportionately large number of species in our morphometric dataset. For instance, Passeriformes, despite being a particularly species-rich clade, is represented by only three species in our dataset. Their small body size could have contributed to their limited representation in skeletal collections and/or quality of specimens. Despite the limitation, our datasets has a good phylogenetic coverage, with three major clades of crown birds (Palaeognathae, Galloanserae, and Neoaves) and a total of 21 orders represented. The other group characterized by long necks, the sauropodomorphs, may be subject to an underrepresentation of long-necked species: larger sauropods are usually known from more fragmentary materials (thereby ineligible for inclusion in this study) [63]. While the distribution of sauropodomorph proportion is not well represented in available data, a rough range is probably captured.

### Phylogenetic analysis of lengths

The evolution of ln 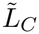 is slightly better explained by a Brownian motion (BM) model (AICc = 20.61) compared to an Ornstein–Uhlenbeck (OU) model (AICc = 23.11). Fitting of two models gave the same estimates of *z*_0_ and *Σ*^2^ (Fig. 1C). Analysis of ln 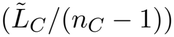 showed similar result: BM (AICc = 10.95) is slightly more supported than OU (AICc = 13.45), but fitting of both models gave the same *z*_0_ and *Σ*^2^ estimates (Fig. 1E). The *z*_0_ estimates together correspond to *n*_*C*_ = 15.30 (Fig. 1D). Analysis of another dataset of bird cervical vertebrae counts [9] showed similar results: BM is explains the divergence of ln (*n*_*C*_ *−* 1) slightly better than OU (AICc = *−*133.4 and AICc = *−*131.3, respectively), and fitting of both models to gave the same estimates of The *z*_0_ and *Σ*^2^. The *z*_0_ estimate corresponds to *n*_*C*_ = 14.67, which is close to that obtained using our morphometric dataset.

### Limitation of relevant datasets

There are two existing datasets that contain cervical phenotypes of diverse bird species but turn out inadequate or unsuitable for the purpose of this study. Böhmer et al. (2019) reported absolute neck lengths and cervical vertebrae counts of about 100 bird species [9]. The neck lengths, however, were not accompanied by lengths of thoracic and/or sacral regions, and cannot be normalized in the same way as we did in this study. Böhmer et al. also reported femur lengths and body masses, which can potentially be used for normalization the neck length; normalization using these variables, however, would only be suitable for comparisons among birds (and perhaps their close relatives), but among diverse groups with more drastically different body plans. Nonetheless, an analysis cervical vertebrae counts reported in this study suggests our morphometric dataset has a matching level of phylogenetic representativeness (see above).

Cerbus et al. (2024) complied a dataset that includes cervical, thoracic, lumbar, sacral, and caudal vertebrae counts of more than 400 extant tetrapod species, including more than 100 birds [16]. Such a dataset could have been usable for a phylogenetic analysis of vertebrae counts, this dataset turns out unusable for the purpose of this study because Cerbus et al. defined cervical vertebrae as vertebrae between the skull and the first rib-bearing vertebrae, ignoring developmental identity as well as other anatomical featured. As a result this practice, vertebrae with a cervical developmental identity but bear rib-like structures could be mis-classified as thoracic. For instance, the two crocodilian species included in the dataset, the American alligator (*Alligator mississippiensis*) and the Siamese crocodile (*Crocodylus siamensis*), were reported to have only two cervical vertebrae [16], presumably because of the presence of cervical ribs in crocodilians; other lines of evidence instead suggest that crocodilians have nine cervical vertebrae [8, 64]. Similarly, cervical vertebrae counts of squamates reported by Cerbus et al. are generally low (no more than five, with most species reported to have two or three) and show inconsistency with previously reported cervical counts of squamates (e.g., [65–67]). While birds may not be substantially affected by the same problem, comparison with other groups can be severely confounded, rendering this dataset unusable for this study.

## Supplementary Tables and Figures

**Table S1:**
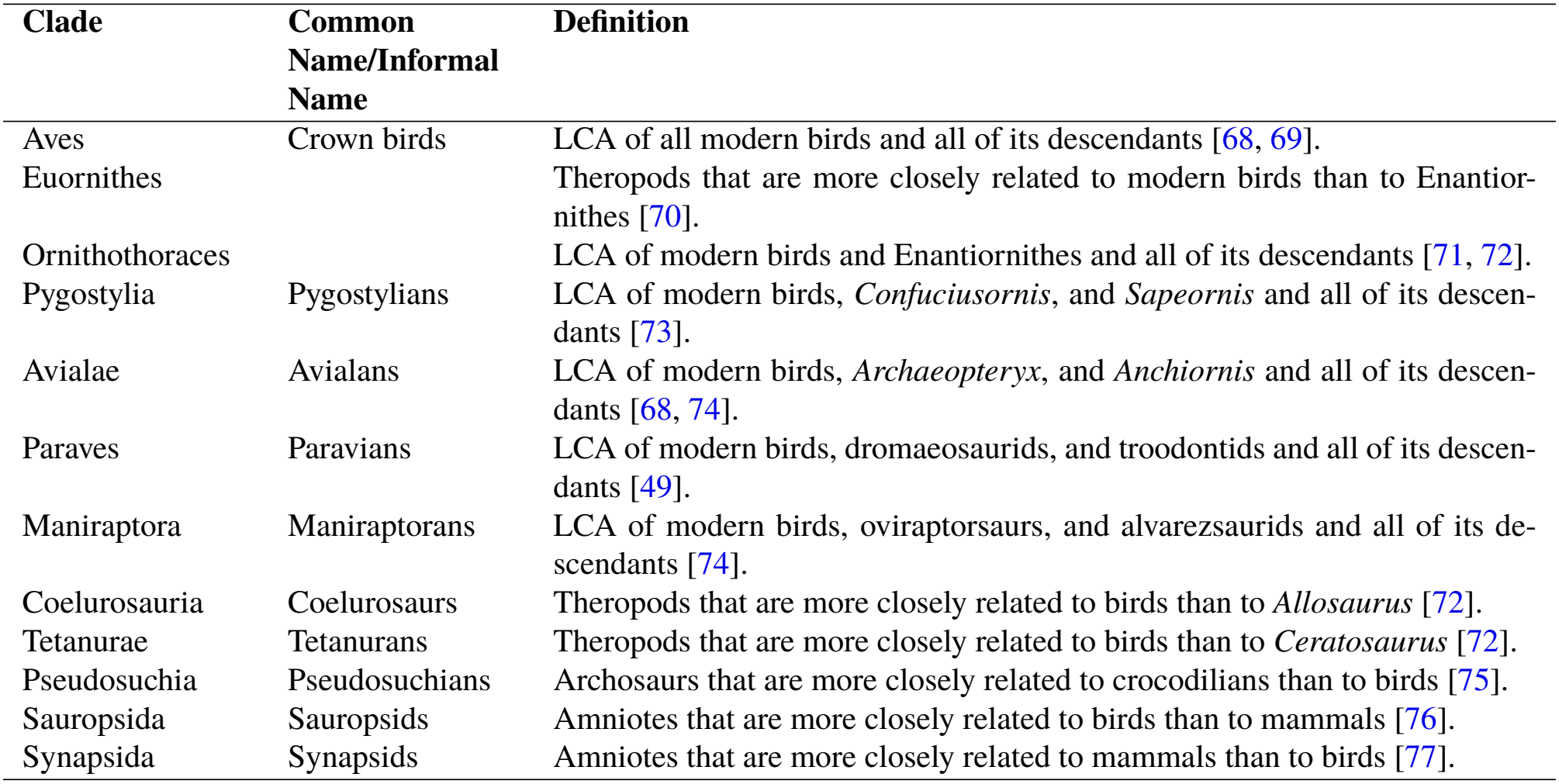
Operational definitions of clades referred to in this study.

| **Clade** | **Common Name/Informal Name** | **Definition** |
|---|---|---|
| Aves | Crown birds | LCA of all modern birds and all of its descendants [68, 69]. |
| Euornithes |  | Theropods that are more closely related to modern birds than to Enantior-nithes [70]. |
| Ornithothoraces |  | LCA of modern birds and Enantiornithes and all of its descendants [71, 72]. |
| Pygostylia | Pygostylians | LCA of modern birds, *Confuciusornis*, and *Sapeornis* and all of its descen-dants [73]. |
| Avialae | Avialans | LCA of modern birds, *Archaeopteryx*, and *Anchiornis* and all of its descen-dants [68, 74]. |
| Paraves | Paravians | LCA of modern birds, dromaeosaurids, and troodontids and all of its descen-dants [49]. |
| Maniraptora | Maniraptorans | LCA of modern birds, oviraptorsaurs, and alvarezsaurids and all of its de-scendants [74]. |
| Coelurosauria | Coelurosaurs | Theropods that are more closely related to birds than to *Allosaurus* [72]. |
| Tetanurae | Tetanurans | Theropods that are more closely related to birds than to *Ceratosaurus* [72]. |
| Pseudosuchia | Pseudosuchians | Archosaurs that are more closely related to crocodilians than to birds [75]. |
| Sauropsida | Sauropsids | Amniotes that are more closely related to birds than to mammals [76]. |
| Synapsida | Synapsids | Amniotes that are more closely related to mammals than to birds [77]. |

**Figure S1:**
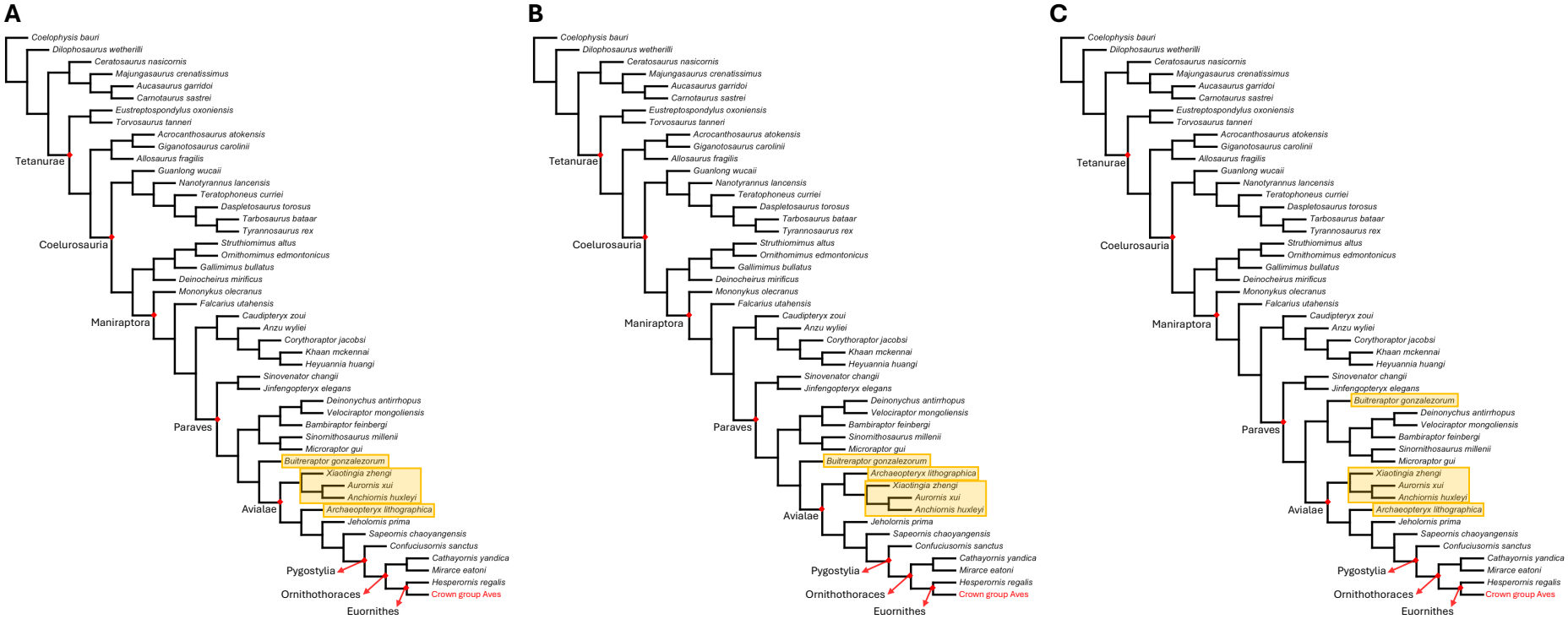
Topologies of theropod trees used in this study. Taxa whose with different placements in different topologies are highlighted. (A) Equal weighting tree from Motta et al. (2025) [49]. (B) Extended implied weighting tree from Motta et al. (2025) [49]. (C) Tree from Pei et al. (2020) [41]. Ancestral nodes of clades referred to in Fig. 2B are shown in each panel.

**Figure S2:**
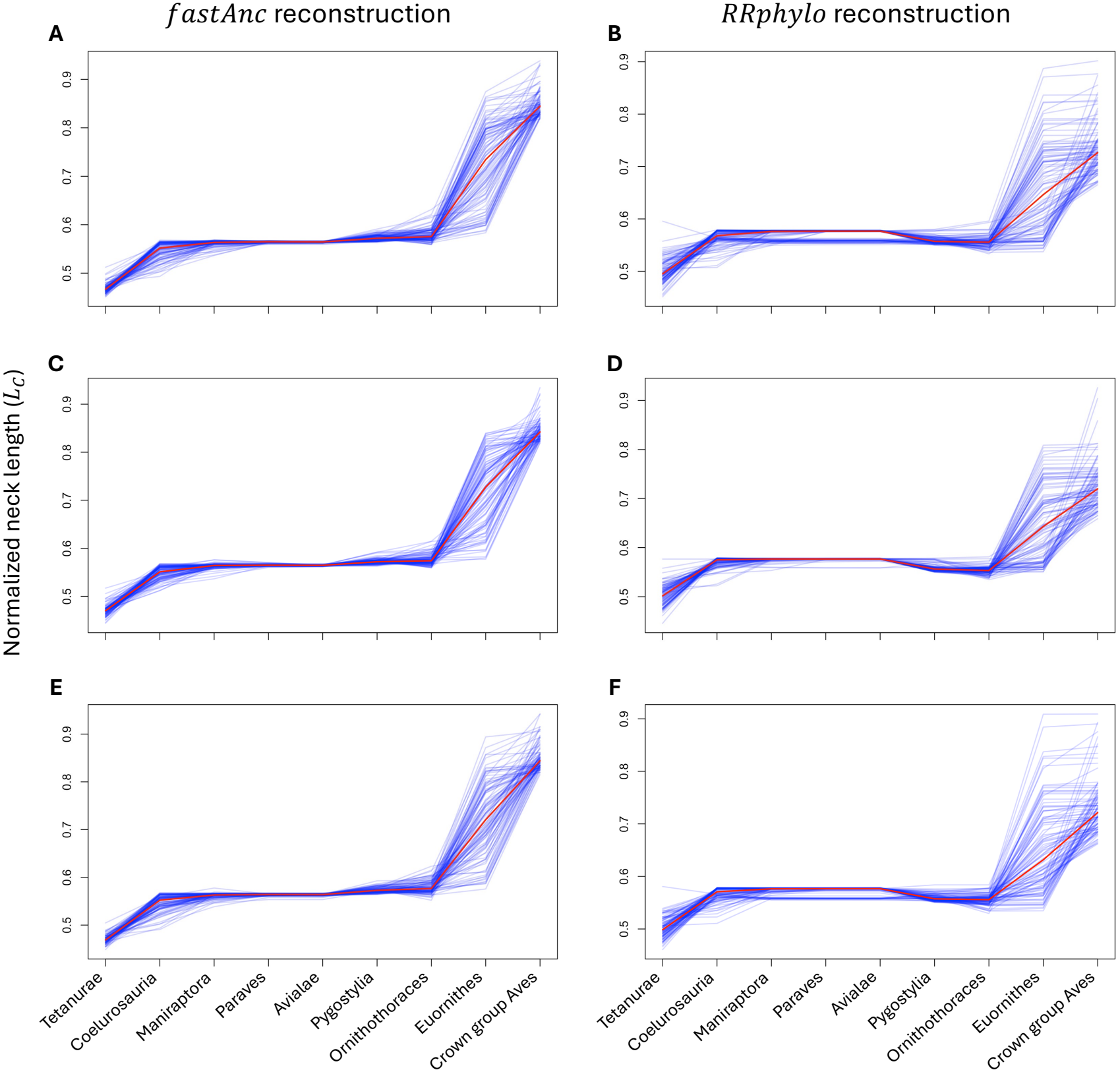
Inferred ancestral states of 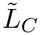 of clades that include crown birds. Notations follow Fig. 2B. (A-B) Inference based on topology in Fig. S1A. (C-D) Inference based on topology in Fig. S1B. (E-F) Inference based on topology in Fig. S1C. (A), (C), and (E): inference using `fastAnc` of `phytools`. (B), (D), and (F): inference using `RRphylo`.

**Figure S3:**
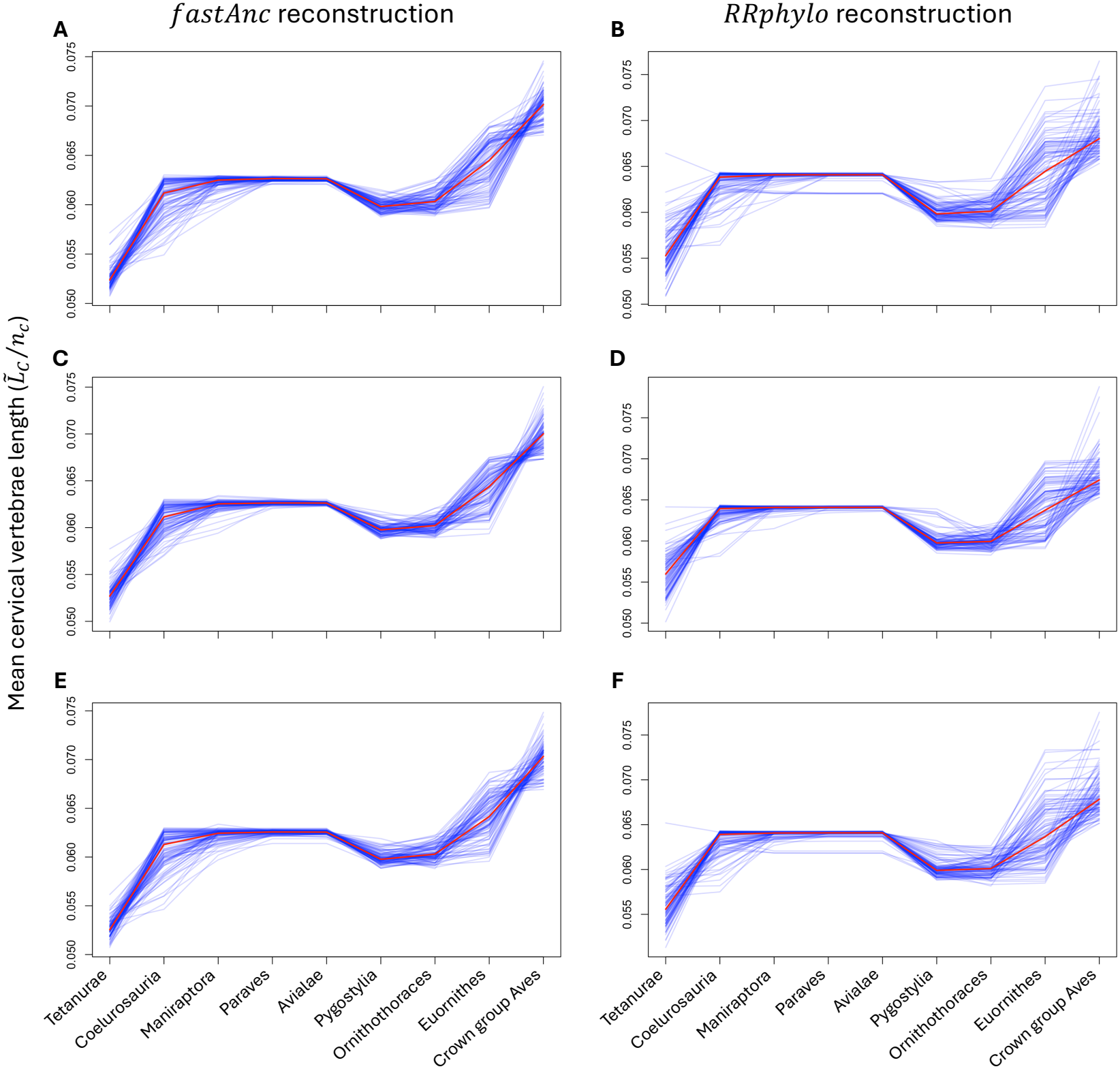
Inferred ancestral states of 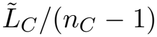 of clades that include crown birds. Notations follow Fig. 2B. (A-B) Inference based on topology in Fig. S1A. (C-D) Inference based on topology in Fig. S1B. (E-F) Inference based on topology in Fig. S1C. (A), (C), and (E): inference using fastAnc of phytools. (B), (D), and (F): inference using `RRphylo`.

**Figure S4:**
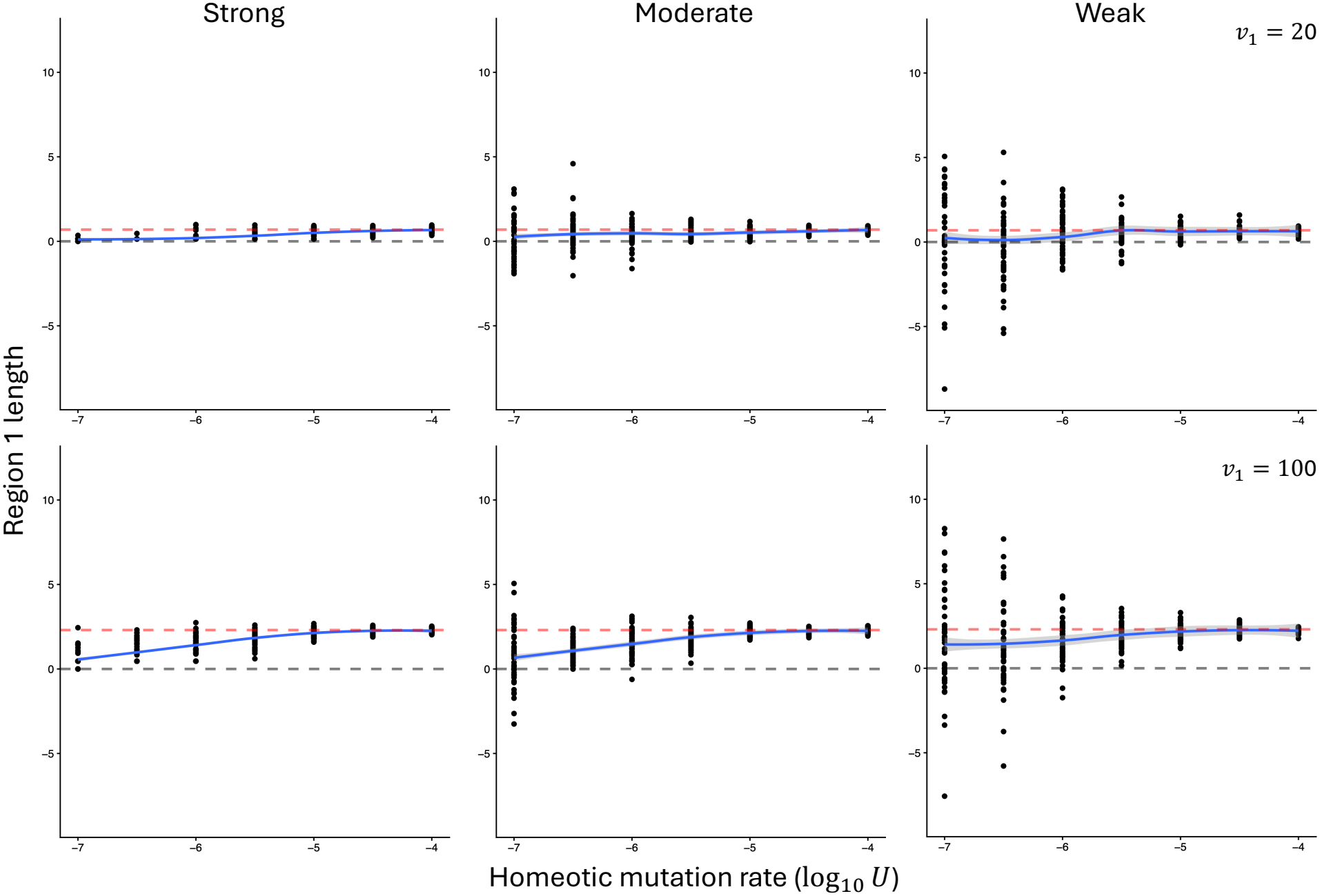
Response to selection on the total length of region 1. Top and bottom rows correspond to scenarios where the optimal region 1 length (*v*_1_) is 2 and 10 times of the ancestral value, respectively. Left, middle, and right columns correspond to scenarios of strong, moderate, and weak stabilizing selection on individual segment length. Notations are otherwise the same as Fig. 3C.

**Figure S5:**
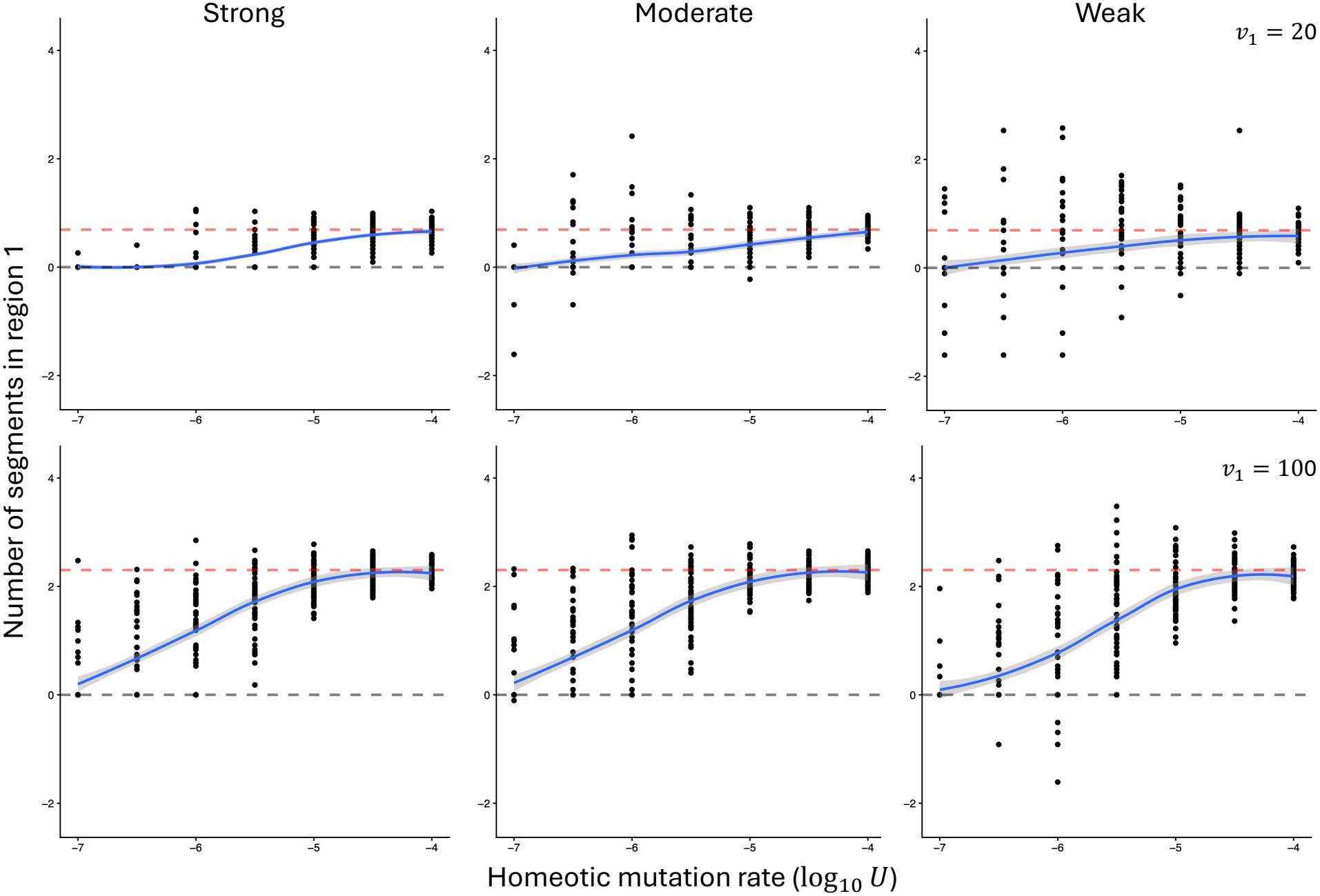
Response of region 1 segment count to selection on the total length of region 1. Top and bottom rows correspond to scenarios where the optimal region 1 length (*v*_1_) is 2 and 10 times of the ancestral value, respectively. Left, middle, and right columns correspond to scenarios of strong, moderate, and weak stabilizing selection on individual segment length. Notations are otherwise the same as Fig. 3D.

**Figure S6:**
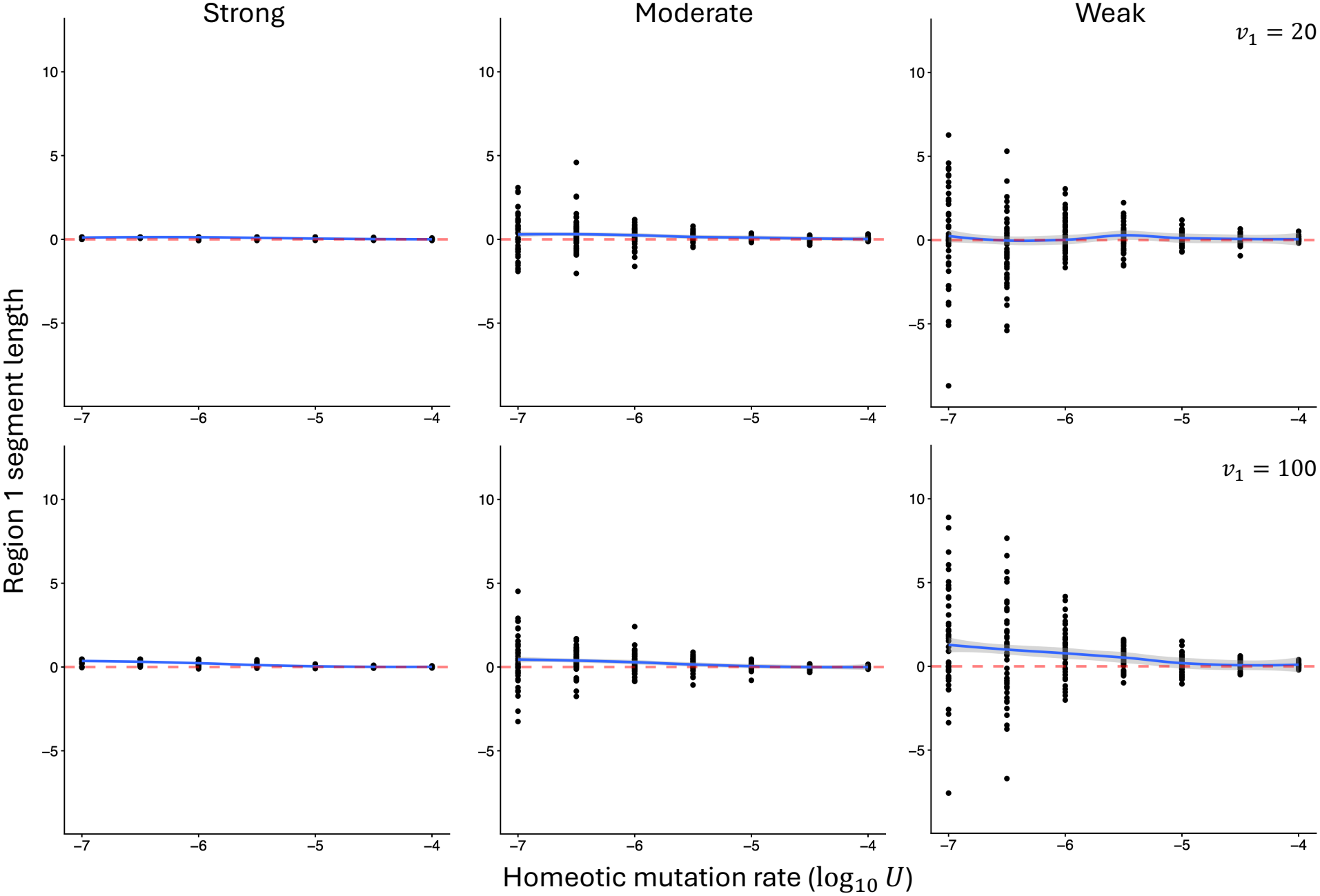
Response of region 1 segment length to selection on the total length of region 1. Top and bottom rows correspond to scenarios where the optimal region 1 length (*v*_1_) is 2 and 10 times of the ancestral value, respectively. Left, middle, and right columns correspond to scenarios of strong, moderate, and weak stabilizing selection on individual segment length. Notations are otherwise the same as Fig. 3E.

